# Rapid, parallel identification of pathways for catabolism of lignin-derived aromatic compounds

**DOI:** 10.1101/160747

**Authors:** Jacob H. Cecil, Joshua K. Michener

## Abstract

This manuscript has been authored by UT-Battelle, LLC under Contract No. DE-AC05-00OR22725 with the U.S. Department of Energy. The United States Government retains and the publisher, by accepting the article for publication, acknowledges that the United States Government retains a non-exclusive, paid-up, irrevocable, world-wide license to publish or reproduce the published form of this manuscript, or allow others to do so, for United States Government purposes. The Department of Energy will provide public access to these results of federally sponsored research in accordance with the DOE Public Access Plan (http://energy.gov/downloads/doe-public-access-plan).

## Introduction

Transposon mutagenesis is a powerful technique in microbial genetics for the identification of genes in uncharacterized pathways^1–3^. However, transposon library screening does not scale well, requiring a new screen for each substrate and extensive oversampling to completely identify all the genes in a given metabolic pathway. A recent technique, dubbed BarSeq^4^, overcomes these limitations by introducing random 20-nucleotide barcodes into each transposon mutant, which can then be tracked using high-throughput DNA sequencing to quantitatively measure the fitness effect of each knockout. Here we show that, when applied to catabolic pathways, barcoded transposon libraries can be used to distinguish redundant pathways, decompose complex pathways into substituent modules, discriminate between enzyme homologs, and rapidly identify previously-hypothetical enzymes in an unbiased genome-scale search. We use this technique to demonstrate that two genes, which we name *desC* and *desD*, are involved in the degradation of the lignin-derived aromatic compound sinapic acid in the non-model organism *Novosphingobium aromaticivorans*. This approach will be particularly useful in the identification of complete pathways suitable for heterologous expression in metabolic engineering.

Microbes are capable of degrading a vast array of natural and non-natural small molecules using an intricate network of metabolic pathways. The characterization of these networks is often the work of many years, particularly for ancestral pathways that can be scattered throughout the genome^5^. Closing a pathway by identifying the entire catalytic ensemble is particularly challenging using random mutagenesis methods, since a search for the last enzyme in a pathway will frequently re-isolate previously-identified enzymes. This challenge is particularly acute when the pathways are intended for use in metabolic engineering, since any missing enzyme renders the entire pathway ineffective^6^. Pathways for the degradation of lignin-derived aromatic compounds provide a clear example of this problem, as these complex pathways are important targets for heterologous expression in industrial microbes and yet rarely found in nature as complete operons^5,7–9^.

We have addressed this problem of catabolic pathway identification using randomly-barcoded transposon mutagenesis (Figure 1)^4^. Using this approach, an entire library of transposon mutants can be screened against a panel of substrates in a single experiment. We chose *Novosphingobium aromaticivorans* DSM12444 (hereafter ‘DSM12444’), a bacterium isolated from contaminated sediment, as a model system for pathway identification^10,11^. This strain has been shown to catabolize a wide range of aromatic compounds, and its catalytic repertoire is predicted to include enzymes that degrade model lignin-derived biaryls such as guaiacylglycerol-β-guaiacyl ether^12^. To begin identifying the genes involved in aromatic degradation in this strain, we constructed a barcoded transposon library in DSM12444. A single round of transposon insertion sequencing was performed to map barcodes to insertion sites, identifying a total of 43,270 unique barcoded insertions in 3117 out of 3897 protein-coding genes. The pooled transposon library was then grown, in triplicate, in minimal medium containing protocatechuate (PCA), 4-hydroxybenzoate (4-HB), coumarate, vanillate, ferulate, syringate, or sinapate as the sole source of carbon and energy. The mixed barcodes were amplified from the inoculum and the final cultures and then sequenced to determine the fitness effect of knocking out each identified gene under the relevant conditions.

**Figure 1:**
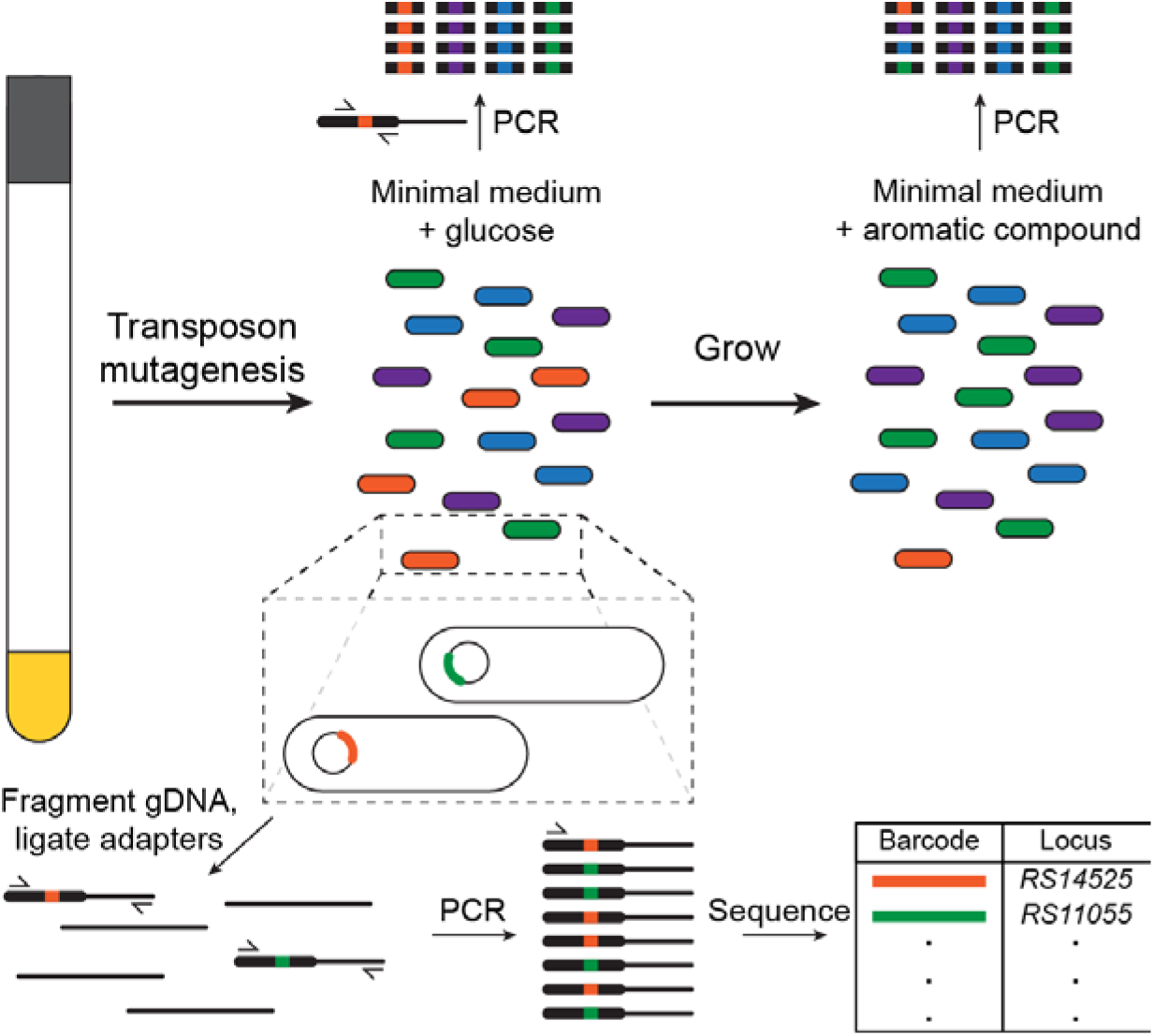
Randomly barcoded transposon mutagenesis allows high-throughput pathway identification. A pool of barcoded transposons is introduced into the target bacterium. A single round of TnSeq maps barcodes (colored inserts) to the insertion locus. The barcodes can be queried through PCR and high-throughput sequencing to determine changes in the population ratio during growth with a specific compound.

Sphingomonads are known to catabolize PCA through 4,5-cleavage of the aromatic ring. DSM12444 contains chromosomal copies of the genes necessary for this cleavage pathway, *ligABCIUJK*, as well as plasmid-encoded genes for the lower 2,3-cleavage pathway for catechol, *xylEGHIJKQ*. During growth with PCA, disrupting either pathway produces a consistent decrease in fitness, averaging a 76% decrease in fitness for an insertion in the *lig* pathway or a 39% decrease in fitness for an insertion in the *xyl* pathway (Figure 2). Replica plating can obscure subtle differences in growth rate resulting from disruption of redundant pathways, but these differences are readily observable in our quantitative fitness measurements. We also note that disruption of the genes encoding ring-cleaving dioxygenases, *ligAB* and *xylE*, do not demonstrate measurable fitness defects. DSM12444 contains at least seven ring-cleaving dioxygenases^13^, and we hypothesize that, when grown in the presence of high concentrations of PCA, these alternate dioxygenases can substitute for the pathway-specific enzymes.

**Figure 2:**
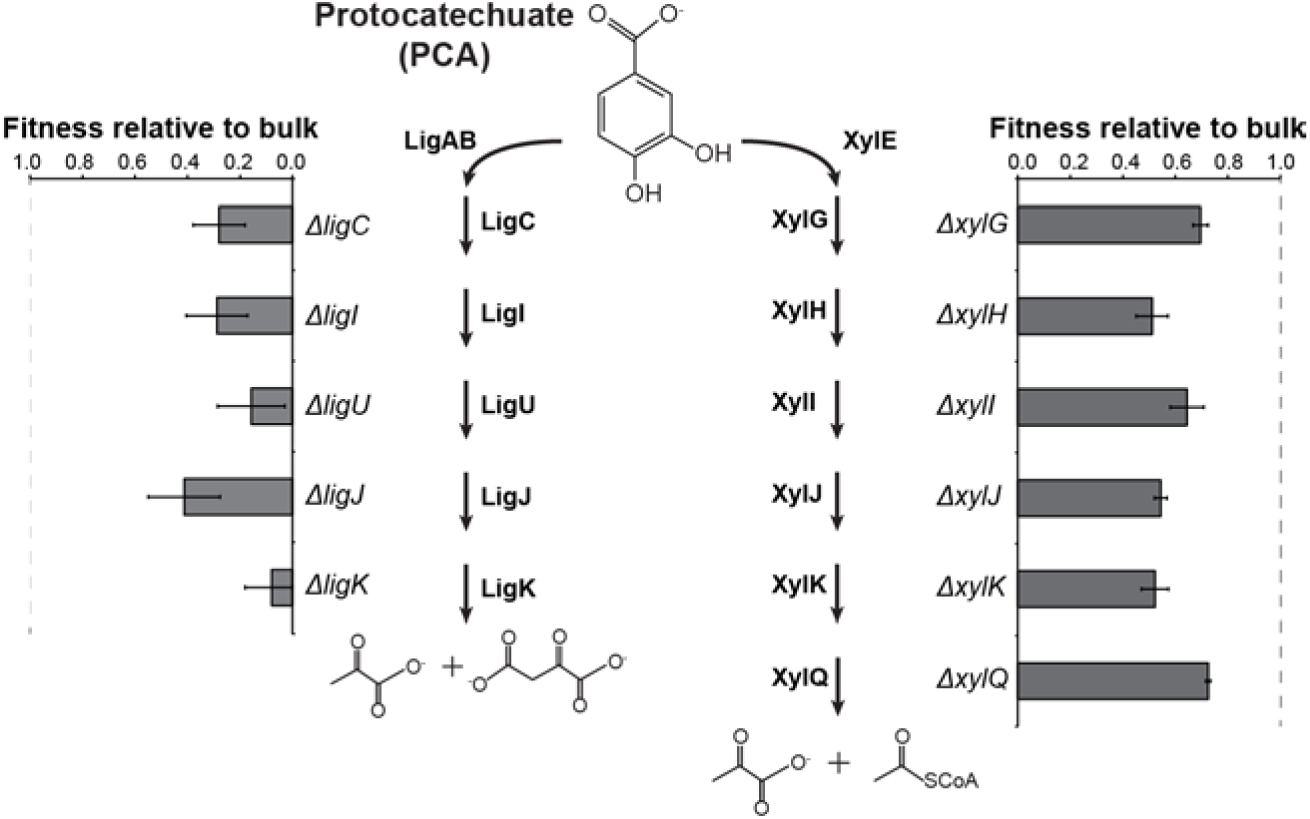
Quantitative fitness measurements identify redundant catabolic pathways for PCA. DSM12444 has two redundant pathways for catabolism of PCA, a 4,5-cleavage pathway and a 2,3-cleavage pathway. Fitness costs for each gene disruption were calculated based on the change in population ratio of every mutant with a transposon insertion in the appropriate gene, and are reported as the competitive fitness relative to the bulk population. Disruption of the canonical 4,5-cleavage pathway produces an average 76% decrease in fitness, while disruption of the 2,3-cleavage pathway decreases fitness by an average of 39%. Error bars show one standard deviation, calculated from three biological replicates.

We next examined the genes involved in degradation of ferulate and vanillate. By measuring the fitness effects during growth not just with the ultimate substrate, ferulate, but also during growth with the intermediates vanillate and PCA, we can decompose the entire pathway for ferulate degradation into modules for the conversion of ferulate to vanillate, vanillate to PCA, and PCA to central metabolites (Figure 3). For example, it is clear from our data that *ferAB* and *ligV* are needed for conversion of ferulate to vanillate, but not for further metabolism of vanillate.

**Figure 3:**
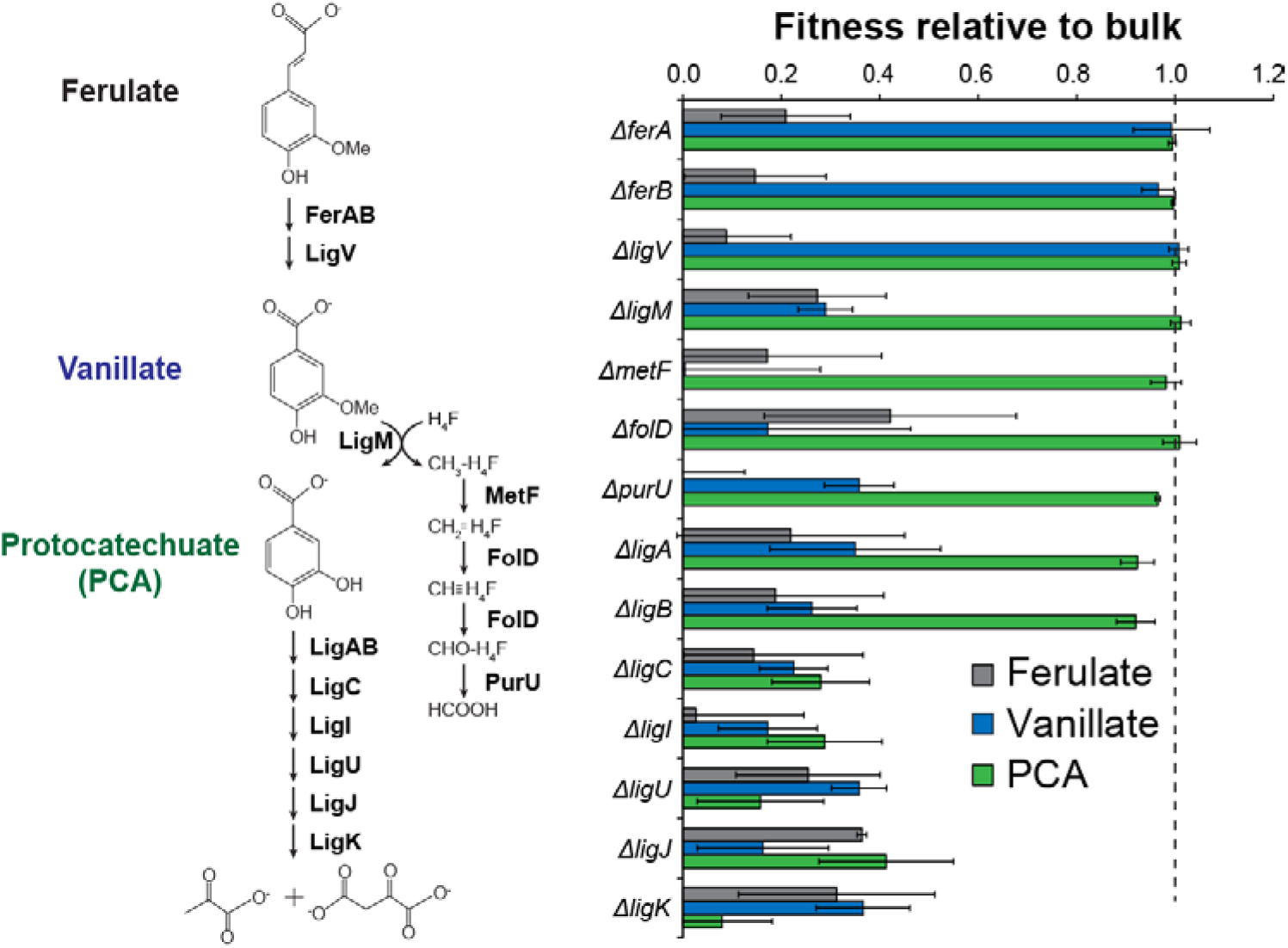
BarSeq measurements enable pathway decomposition. Fitness values were measured during growth with ferulate, vanillate, and PCA, and compared to the bulk fitness. Comparison of the fitness values with different substrates makes the ordering of genes in the pathway readily apparent. Error bars show one standard deviation, calculated from three biological replicates.

Decomposing a pathway in this manner will simplify the process of matching unknown enzymes to their catalytic activities. However, and in contrast to growth with PCA, growth with vanillate or ferulate requires the genes for the PCA ring-cleaving dioxygenase, *ligAB*. When PCA is an intermediate rather than a substrate, its concentration is likely to be lower, which in turn makes the PCA-specific ring-cleaving dioxygenase more critical. In general, we find that the genetic requirements are more stringent for growth with an upper pathway substrate than an intermediate, and this effect could complicate efforts at pathway elucidation. Prior knowledge of the genes involved in ferulate catabolism^14^, accumulated over many years’ of work, certainly aided in identifying relevant genes. However, this knowledge was not strictly necessary. For example, eleven genes are shown in Figure 3 as being involved in vanillate degradation. Disruption of these genes produced eleven of the top twelve fitness defects during growth with vanillate, and the twelfth is a hypothetical gene with homology to DNA repair restriction endonucleases.

Previous work in *Sphingobium* sp. SYK-6 has demonstrated that the vanillate demethylase LigM transfers a methyl group from the aromatic substrate to tetrahydrofolate. Pathways for conversion of methyltetrahydrofolate to tetrahydrofolate and formate have been characterized in a variety of organisms, involving sequential oxidations by MetF, FolD, and PurU. However, many organisms, including DSM12444, contain multiple homologs of these genes, and the specific catabolic homologs may be necessary for proper function of a heterologously-expressed pathway^15^. During growth with vanillate, specific *metF* and *folD* homologs are required for growth, while the alternative homologs are entirely dispensable (Supplementary Figure 1). The correct *metF* homolog is located in a gene cluster with *ligM*, consistent with its role in vanillate demethylation. However, the dispensable *folD2* homolog is chromosomal, while the necessary *folD* homolog is located on plasmid pNL2.

Two pathways are known to deacetylate cinnamic acid derivatives to the corresponding benzoic acid derivatives, either through β-oxidative deacetylation to directly yield the acid or through a combined hydratase/lyase that produces a free benzaldehyde, which can be further oxidized to the acid. The latter pathway is present in *Sphingobium* sp. SYK-6, with FerA and FerB performing the deacetylation^16^, followed by oxidation by LigV or DesV^17,18^. Likewise, β-oxidative routes have been described for ferulate and coumarate in *Rhodococcus jostii* RHA1^19^ and for ferulate, coumarate, and caffeate in *Agrobacterium fabrum*^20^. However, no β-oxidative pathway has previously been demonstrated for conversion of sinapate to syringate.

DSM12444 contains homologs of FerA, FerB, and LigV, and disrupting any of the associated genes severely decreases fitness with coumarate, ferulate, and sinapate (Supplementary Figure 2A). However, disruption of an additional three-gene cluster, SARO_RS14535, SARO_RS14545, and SARO_RS14550, also significantly reduces growth with sinapate (Supplementary Figure 2B). SARO_RS14535 and SARO_RS14545 have homology to the phenylhydroxypropionyl-CoA dehydrogenase Atu1415 and the 4-hydroxyphenyl-beta-ketoacyl-CoA hydrolase Atu1421 from *A. fabrum*, respectively^20^. By homology, SARO_RS14550 is likely to be an aromatic aldehyde dehydrogenase, presumably with similar functionality as LigV or DesV ^17^. We hypothesize that these enzymes act in parallel with FerA, FerB, and LigV to deacetylate cinnamic acid derivatives. SARO_RS14535 and SARO_RS14550 appear to be specific to sinapate, while SARO_RS14545 contributes equally to catabolism of ferulate, coumarate, and sinapate. *Sphingobium* sp. SYK-6 contains homologs to SARO_RS14535 and SARO_RS14545, SLG_RS06215 and SLG_RS06210, respectively. No additional CoA-ligases, aside from FerA, were identified in our analysis as being involved in activation of sinapate, ferulate, or coumarate.

DSM12444 grows efficiently with both syringate and sinapate. In *Sphingobium* sp. SYK-6, three pathways have been identified for syringate degradation. The first step in each pathway is demethylation by DesA to produce 3-*O*-methylgallate (3MGA)^21^. A *desA* homolog is present in DSM12444 and required for growth with sinapate, reducing fitness by 91% when disrupted (Supplementary Figure 3). Conversion of 3MGA into 4-oxalomesaconate (OMA) requires two steps, ring-opening and demethylation. In the first pathway, LigM demethylates 3MGA to gallate, followed by oxidation by DesB or LigAB to yield OMA^22–24^. In the second pathway, DesZ or LigAB oxidizes and demethylates 3MGA to 2-pyrone-4,6-dicarboxylate (PDC), followed by hydrolysis by LigI^25^. Finally, the third pathway proceeds through ring opening by LigAB, followed by at least one uncharacterized reaction to yield OMA^26^. No homologs of *desB* or *desZ* are present in DSM12444. Growth with sinapate requires *ligABUJK* but not *ligI*, suggesting that conversion through PDC is not a major route (Supplementary Figure 3).

Since our fitness measurements could efficiently recapitulate known pathways for aromatic catabolism, we sought to use these data to identify the uncharacterized enzymes in this pathway for syringate degradation. Cleavage of 3-*O*-methylgallate produces a mixture of stereoisomers of 4-carboxy-2-hydroxy-6-methoxy-6-oxohexa-2,4-dienoate (CHMOD)^25^. The conversion of mixed isomers of CHMOD into OMA is expected to involve a methylesterase and a cis-trans isomerase. Based on homology, SARO_RS14525 is predicted to encode a methylesterase and SARO_RS14530 a glutathione S-transferase. Glutathione S-transferases can function as cis-trans isomerases, such as in the case of maleylpyruvate isomerase^27^. Disruption of either gene is detrimental during growth with sinapate, while being dispensable during growth with PCA, 4-HB, coumarate, vanillate, and ferulate (Supplementary Figure 4). Accordingly, we propose that SARO_RS14525, which we name *desC*, demethylates CHMOD, and SARO_RS14530, which we name *desD*, isomerizes one of the intermediates. Notably, *Sphingobium* sp. SYK-6 contains homologs of both *desC* and *desD*, and *desC* occurs in a cluster with the homologs of RS14535, RS14545, *ligM, metF*, and *ftfL*.

To confirm these results, we constructed clean deletions of *ligA, ferA, ligM, desA, desC*, and *desD* in DSM12444. The deletion strains were grown in minimal medium containing glucose, 4-HB, ferulate, or sinapate as the sole source of carbon and energy, and their growth rates were compared to the wild type (Figure 4). As expected, none of the mutations affected growth with glucose, and only the *ligA* deletion decreased growth with 4-HB. Deletion of *ligA, ligM*, or *ferA* significantly decreased growth with ferulate, while deletion of *desA, desC*, or *desD* had no effect. In contrast, deletion of *ligM* had no impact on growth with sinapate, while deletion of *desA* and *desC* severely decreased growth and deletion of *desD* had a moderate effect. Since *ligM* is dispensable but *desC* is required for growth with sinapate, we conclude that DSM12444 degrades sinapate solely through CHMOD. We hypothesize that, as shown in SYK-6, cleavage of the 3-MGA ring by LigAB produces a mixture of stereoisomers, which are resolved by DesD. In the absence of *desD*, growth is permitted, but inefficient.

**Figure 4:**
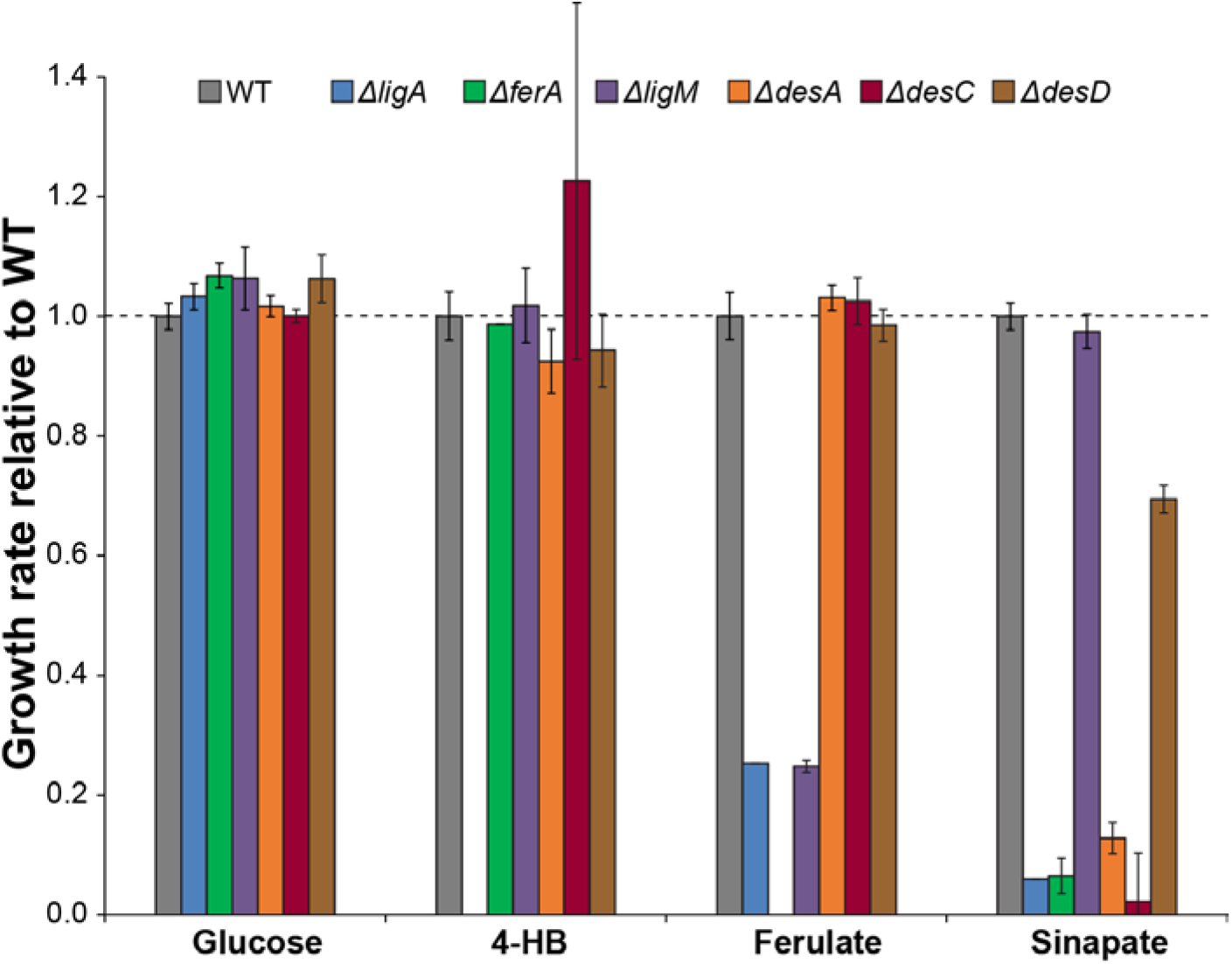
Gene deletions confirm bulk fitness measurements and identify enzymes for sinapate degradation. Mutant strains with clean deletions of the indicated genes were grown in minimal medium containing the indicated carbon source. Disruptions of the newly-identified genes *desC* and *desD* decrease growth with sinapate, but not with any of the other substrates tested. Error bars show one standard deviation, calculated from three biological replicates.

Due to the inherent heterogeneity of lignin, its degradation produces a wide range of aromatic compounds^9^. Engineering microbes to productively convert these aromatic compounds into valuable fuels and chemicals will require the identification, characterization, and integration of multiple, complex metabolic pathways. To meet this challenge, the tools employed must scale with the scope of the networks^6^. In this work, we have shown that randomly-barcoded transposon mutagenesis provides the necessary scale for pathway discovery, by rapidly and quantitatively identifying genes that affect catabolism across panels of substrates.

## Materials and Methods

### Media and chemicals

All chemicals were purchased from Sigma-Aldrich (St. Louis, MO) or Fisher Scientific (Fairlawn, NJ) and were molecular grade. All oligonucleotides were ordered from IDT (Coralville, IA). *E. coli* were routinely cultivated at 37 °C in LB, while *N. aromaticivorans* were grown at 30 °C in LB or DSM Medium 457 containing the indicated carbon source. Aromatic monomers were dissolved in DMSO at 100 g/L and added at a working concentration of 1 g/L. As needed, cultures were supplemented with kanamycin at 50 mg/L, streptomycin at 100 mg/L, or diaminopimelic acid (DAP) at 60 mg/L.

### Strains and plasmids

*N. aromaticivorans* DSM12444 was purchased from the DSMZ (Braunschweig. Germany). Strain APA766, comprising the conjugation donor WM3064 containing the barcoded Tn5 vector pKMW7, was a gift from Adam Deutschbauer. The parental deletion plasmid, pAK405, was a gift from Julia Vorholt (Addgene plasmid # 37114). Deletion plasmids pJM269-280 were synthesized *de novo* and cloned into pAK405 by Genscript (Piscataway, NJ).

### Transposon library construction and indexing

DSM12444 and APA766 were grown overnight in LB, containing kanamycin and DAP as appropriate. DSM12444 was then diluted 1:100 into 50 mL of fresh LB and regrown to mid-log phase. Both cultures were pelleted by centrifugation, mixed at a ratio of 5:1 recipient:donor, and plated on sterile filter paper placed on an LB+DAP agar plate. After overnight incubation at 30 °C, the filter paper was removed, vortexed with 5 mL of fresh LB, pelleted by centrifugation, resuspended in 1 mL of fresh LB, and plated on 20 LB+kanamycin plates. Colonies were observed after 3 days of incubation at 30 °C. The colonies were scraped from the plates, combined into a single mixed culture, and grown overnight in 50 mL of LB+kanamycin at 30 °C. Aliquots of the combined library were stored in 7% DMSO at -80 °C for further analysis. Additionally, genomic DNA was prepared using a DNeasy Blood and Tissue kit (Qiagen, Valencia, CA) according to the manufacturer’s directions.

Barcode indexing was performed based on published protocols^4,28^. Briefly, genomic DNA was fragmented by sonication using a Bioruptor Plus (Diagenode, Ougrée, Belgium) and size-selected to approximately 300bp using Agencourt AMPure XP beads (Beckman Coulter, Indianapolis, IN). Size selection was confirmed using a Bioanalyzer DNA1000 chip (Agilent, Santa Clara, CA). End repair, dA-tailing, and adaptor ligation were performed using the NEBNext kit according to the manufacturer’s directions (NEB, Ipswitch, MA). Custom adaptors were synthesized by IDT following published sequences^4^. In lieu of a final size selection, two sequential purifications using AMPure XP beads were performed instead^28^. The final transposon enrichment PCR was performed using KAPA HiFi HotStart ReadyMix^28^ using custom indexing and transposon-specific primers^4^. The library was analyzed using a DNA1000 chip, quantified using a Qubit fluorimeter (ThermoFisher, Waltham, MA), and sequenced on an Illumina MiSeq, using v2 chemistry with paired-end 150bp reads (Illumina, San Diego, CA). Reads were analyzed using custom scripts as described previously^4^.

### Pooled fitness measurements

An aliquot of the DSM12444 library was thawed, diluted into 50mL of LB+kanamycin, and grown overnight at 30 °C. The culture was then diluted 1:100 into three flasks, each containing MM457+kanamycin+2 g/L glucose and grown to saturation. Time-zero samples were taken from each flask, and each culture was then diluted to an optical density of 0.005 in 5mL of MM457+kanamycin containing the appropriate carbon sources. Cultures were grown to saturation at 30 °C and then frozen for later analysis. Genomic DNA was isolated and barcodes were amplified using custom indexing primers as described previously ^4^. Barcode amplicons were quantified using a Qubit fluorimeter, pooled, and sequenced at the Joint Genome Institute on a HiSeq 2500 lane. Barcode frequencies and gene fitness values were calculated using custom scripts as described previously^4^.

### Gene deletions

In-frame gene deletions were constructed using a kanamycin/*rpsL* suicide vector^29^. Briefly, appropriate deletion cassettes were synthesized and cloned into pAK405. The resulting plasmids were transformed into strain WM6026, a conjugation-proficient DAP auxotroph. Plasmids were transferred to DSM12444 by conjugation following the same protocol as for library construction. After selecting for single crossovers on LB+kanamycin plates, colonies were patched on LB+kanamycin to confirm plasmid integration, grown in the absence of selection, and then plated on LB+streptomycin plates. Colonies were patched on LB+streptomycin and replica plated on LB+streptomycin+kanamycin to confirm double recombination. Deletions were identified by colony PCR using locus-specific primers and confirmed by Sanger sequencing of the amplicons.

### Growth rate measurements

Strains were grown to saturation overnight in minimal medium with 2 g/L glucose. They were then diluted 100x into fresh medium containing the appropriate carbon source and grown as triplicate 100 µL cultures in a Bioscreen C plate reader (Oy Growth Curves Ab Ltd, Helsinki, Finland). Growth was monitored using optical density at 600 nm (OD_600_). Growth rates were calculated during exponential growth using CurveFitter software^30^.

## Acknowledgements

Strains for barcoded transposon library construction were provided by Adam Deutschbauer at the Lawrence Berkeley National Laboratory. DNA shearing was performed at the UT-Knoxville Genomics Core, with assistance from Joe May and Veronica Brown. Dawn Klingeman performed the index sequencing reaction. Barcode sequencing was performed by Christa Pennacchio and Matthew Blow at the Joint Genome Institute. The work conducted by the U.S. Department of Energy Joint Genome Institute, a DOE Office of Science User Facility, is supported by the Office of Science of the U.S. Department of Energy under Contract No. DE-AC02-05CH11231. Oak Ridge National Laboratory is managed by UT-Battelle, LLC, for the DOE under Contract No. DE-AC05-00OR22725.

## Author Contributions

JKM conceived of the project. JC and JKM designed and performed experiments, analyzed the data, and wrote the paper.

